# Phylogenomic analyses shed light on the relationships of chiton superfamilies and shell-eye evolution

**DOI:** 10.1101/2022.12.12.520088

**Authors:** Xu Liu, Julia D. Sigwart, Jin Sun

**Author notes:** Corresponding author, Jin Sun or Julia D. Sigwart.

## Abstract

Mollusca is the second-largest animal phylum with over 100,000 species among eight distinct taxonomic classes. Across 1000 living species in the class Polyplacophora, chitons have a relatively constrained morphology but with some notable deviations. Several genera possess “shell eyes”, true eyes with a lens and retina that are embedded within the dorsal shells, which represent the most recent evolution of animal eyes. The phylogeny of major chiton clades is mostly well established, in a set of superfamily and higher-level taxa supported by various approaches including multiple gene markers, mitogenome-phylogeny and phylotranscritomic approaches as well as morphological studies. However, one critical lineage has remained unclear: *Schizochiton* was controversially suggested as a potential independent origin of chiton shell eyes. Here, with the draft genome sequencing of *Schizochiton incisus* (superfamily Schizochitonoidea) plus assembly of transcriptome data from other polyplacophorans, we present phylogenetic reconstructions using both mitochondrial genomes and phylogenomic approaches with multiple methods. Phylogenetic trees from mitogenomic data are inconsistent, reflecting larger scale confounding factors in molluscan mitogenomes. A consistent robust topology was generated with protein coding genes using different models and methods. Our results support Schizochitonoidea is a sister group to other Chitonoidea in Chitonina, in agreement with established classification. This suggests that the earliest origin of shell eyes is in Schizochitonoidea, which were also gained secondarily in other genera in Chitonoidea. Our results have generated a holistic review of the internal relationship within Polyplacophora, and a better understanding on the evolution of Polyplacophora.

## Introduction

Molluscs represent the second most species rich animal phylum with the broadest morphological disparity of body plans. The class Polyplacophora, also known as chitons, includes around 1000 living species and over 400 fossil species (Stebbins et al. 2009). Chitons are exclusively marine, and their most distinctive feature is eight separate aragonitic valves or plates on their dorsal side (Ladd 1966; Stebbins et al. 2009; Irisarri et al. 2020). They attach to the substratum with a muscular ventral foot and feed with an iron-mineralised radula (Joester et al. 2016). They have no head or cephalised senses, and therefore lack conventional eyes. However, the dorsal valves are densely innervated with a complex array of sensory pores called aesthetes which can have densities of over 1000 mm^-2^.

Aesthete pores are present in all chitons, with substantial differences in morphology, size, arrangement, densities, and presumably also functions, and aesthete morphology is often used to discriminate species in taxonomic descriptions (Sirenko 2006). A number of chiton species are demonstrably photosensitive, and some have pigmented aesthetes that apparently function as photoreceptors. In the most elaborate variation, in a few genera, some of the larger “megalaesthete” pores have further developed into shell eyes. These are true eyes, embedded in the shell matrix, with a crystalline lens and a pigmented photoreceptive retina (Sigwart et al. 2021).

The evolution of chiton shell eyes occurred much more recently than any other animal eyes. The oldest fossil shell eyes are known from the fossil genus *Incissiochiton* from the lower Palaeocene (61-66 Mya), which is a member of the family Schizochitonidae, the only family in a superfamily Schizochitonoidea (Sirenko 2006; Sirenko 2013). Members of Schizochitonidae (*Incissiochiton* and the Recent genus *Schizochiton*), as well as species in the two subfamilies Acanthopleurinae and Toniciinae, possess shell eyes. The only previous molecular phylogenetic study that included *Schizochiton* dates back to 2003 with five gene fragments (Okusu et al. 2003), and those authors suggested that the phylogenetic position of *S. incisus* in those analyses was “unstable” and deserved further discussion. Most importantly, the unresolved phylogenetic position of *Schizochiton* raised the possibility that shell eye structures evolved not only relatively recently, but in two separate events. However, in the last 20 years this hypothesis has not been tested further, due to a lack of appropriate specimen material for molecular data from this important lineage *Schizochiton*.

Phylogenetic systematics of Polyplacophora has been developed using both morphological and molecular characters (Albano 2021). Extant chitons are divided into three well-resolved orders: Lepidopleurida, Callochitonida, and Chitonida (Giribet et al. 2020). Lepidopleurida consists of mainly deep-sea species with distinctive morphological synapomorphies including aesthete arrangement, gills, and a specialized sense organ called the Schwabe Organ (Sigwart et al. 2014). The position of *Callochiton* was equivocal in earlier studies but usually resolved as siter to Chitonida (Koch et al. 1990; Sigwart et al. 2013) and the single family Callochitonidae is now recognized as comprising a separate order-ranked clade Callochitonida (Sigwart et al. 2013; Giribet et al. 2020; Moles et al. 2021). Most living chitons are in the order Chitonida, which is further divided into two suborders, Chitonina (including two superfamilies, Chitonoidea and Schizochitonoidea) and Acanthochitonina (including two superfamilies, Mopalioidea and Cryptoplacoidea). The backbone phylogeny of chitons is well understood especially at the level of superfamilies, for all clades except for Schizochitonoidea.

Various genomic and transcriptomic data in Polyplacophora are now available on NCBI but were generated independently for several different research purposes (Table 1). There are only two chiton genomes available, *Acanthopleura granulata* (Varney et al. 2021) and *Hanleya hanleyi* (Varney et al. 2022). Meanwhile, two independent phylogenomic studies based on transcriptome sequencing, generated data for species and genera that cover all Recent superfamilies: *Callochiton, Tonicia schrammi, Chiton tuberculatus*, *Chiton marmoratus, Chaetopleura apiculata, Lepidozona mertensii, Mopalia muscosa, Katharina tunicata, Tonicella lineata, Nutallochiton* sp., *Cryptoplax japonica* and *Crytoplax larvaeformis* (Varney et al. 2021) and *Lepidopleurus cajetanus* (SRX5063921), *Callochiton septemvalvis*, *Stenoplax bahamensis*, *Cryptoplax japonica* and *Choneplax lata* (Moles et al. 2021). There are also some other studies examining the gene expression profiles, which includes *Leptochiton cascadiensis* (Halanych et al. 2014), *Acanthopleura loochooana* (Liu et al. 2022), *Rhyssoplax olivacea* (Riesgo et al. 2012), *Cryptochiton stelleri* (Nemoto et al. 2019), *Acanthochitona crinita* (De Oliveira et al. 2016), *Acanthochitona rubrolineata* (SRP179406) and *Acanthochitona fascicularis* (SRR13862580). These data collection can support a phylogenomic construction with larger taxon coverage. And due to the important potion of *Schizochiton* for us to better understand chiton evolution, we newly sequenced and assembled the genome and mitogenome of *Schizochiton incisus*. Combining this with other available chiton data from NCBI and previous studies, we aimed to reconstruct a phylogeny of Polyplacophora at the superfamily level with different phylogenomics inferences and tree reconstruction methods, specifically to test the position of *S. incisus* and Schizochitonoidea.

**Table 1.**
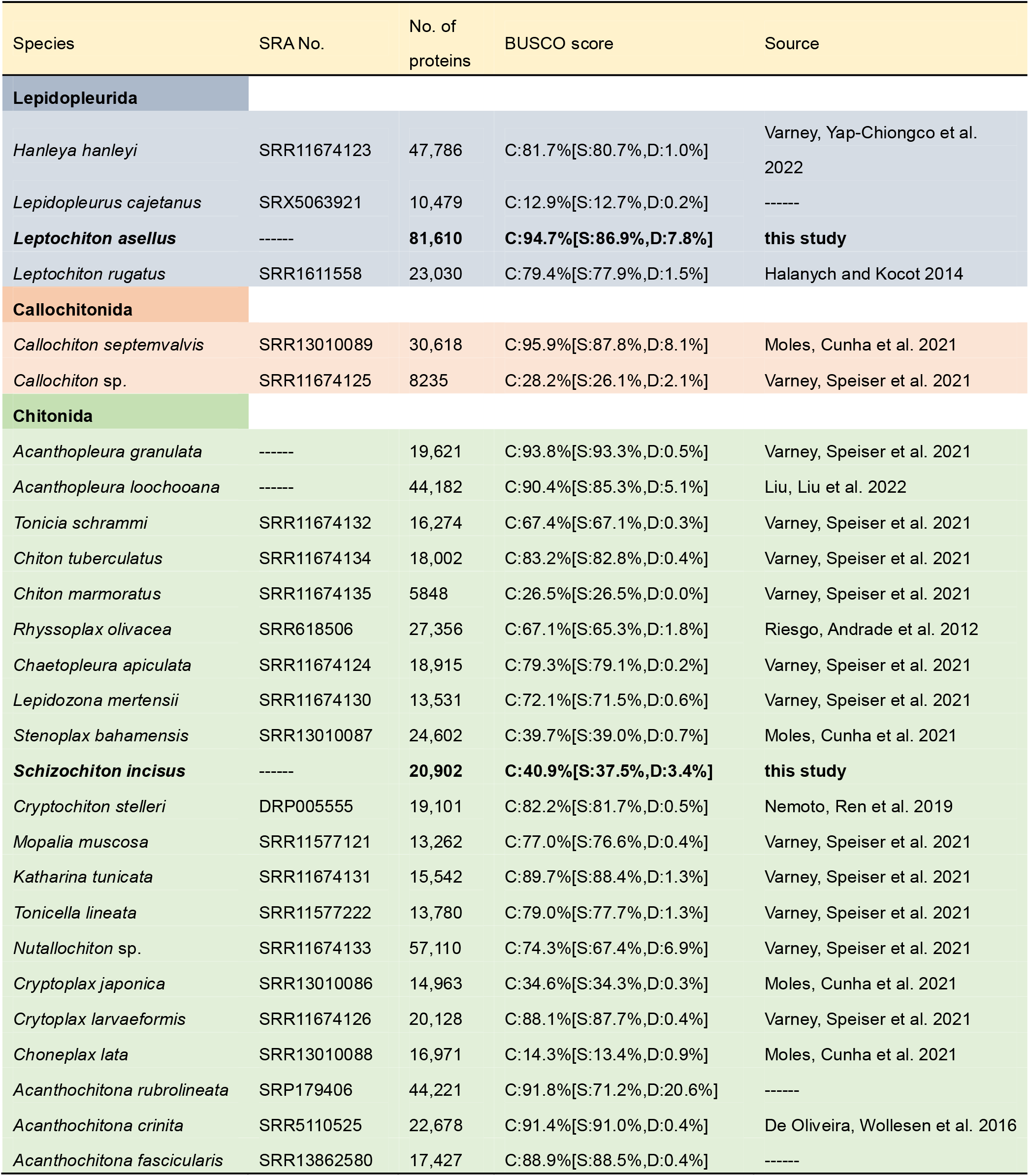
Statistics of chiton genomes and transcriptomes used in this study, including number of contigs and BUSCO scores after filtering.

| Species | SRA No. | No. of proteins | BUSCO score | Source |
| --- | --- | --- | --- | --- |
| **Lepidopleurida** |  |  |  |  |
| *Hanleya hanleyi* | SRR11674123 | 47,786 | C:81.7%[S:80.7%,D:1.0%] | Varney, Yap-Chiongco et al. 2022 |
| *Lepidopleurus cajetanus* | SRX5063921 | 10,479 | C:12.9%[S:12.7%,D:0.2%] | ----- |
| ***Leptochiton asellus*** | ----- | **81,610** | **C:94.7%[S:86.9%,D:7.8%]** | **this study** |
| *Leptochiton rugatus* | SRR1611558 | 23,030 | C:79.4%[S:77.9%,D:1.5%] | Halanych and Kocot 2014 |
| **Callochitonida** |  |  |  |  |
| *Callochiton septemvalvis* | SRR13010089 | 30,618 | C:95.9%[S:87.8%,D:8.1%] | Moles, Cunha et al. 2021 |
| *Callochiton* sp. | SRR11674125 | 8235 | C:28.2%[S:26.1%,D:2.1%] | Varney, Speiser et al. 2021 |
| **Chitonida** |  |  |  |  |
| *Acanthopleura granulata* | ----- | 19,621 | C:93.8%[S:93.3%,D:0.5%] | Varney, Speiser et al. 2021 |
| *Acanthopleura loochooana* | ----- | 44,182 | C:90.4%[S:85.3%,D:5.1%] | Liu, Liu et al. 2022 |
| *Tonica schrammi* | SRR11674132 | 16,274 | C:67.4%[S:67.1%,D:0.3%] | Varney, Speiser et al. 2021 |
| *Chiton tuberculatus* | SRR11674134 | 18,002 | C:83.2%[S:82.8%,D:0.4%] | Varney, Speiser et al. 2021 |
| *Chiton marmoratus* | SRR11674135 | 5848 | C:26.5%[S:26.5%,D:0.0%] | Varney, Speiser et al. 2021 |
| *Rhyssoplax olivacea* | SRR618506 | 27,356 | C:67.1%[S:65.3%,D:1.8%] | Riesgo, Andrade et al. 2012 |
| *Chaetopleura apiculata* | SRR11674124 | 18,915 | C:79.3%[S:79.1%,D:0.2%] | Varney, Speiser et al. 2021 |
| *Lepidozona mertensii* | SRR11674130 | 13,531 | C:72.1%[S:71.5%,D:0.6%] | Varney, Speiser et al. 2021 |
| *Stenoplax bahamensis* | SRR13010087 | 24,602 | C:39.7%[S:39.0%,D:0.7%] | Moles, Cunha et al. 2021 |
| ***Schizochiton incisus*** | ----- | **20,902** | **C:40.9%[S:37.5%,D:3.4%]** | **this study** |
| *Cryptochiton stelleri* | DRP005555 | 19,101 | C:82.2%[S:81.7%,D:0.5%] | Nemoto, Ren et al. 2019 |
| *Mopalia muscosa* | SRR11577121 | 13,262 | C:77.0%[S:76.6%,D:0.4%] | Varney, Speiser et al. 2021 |
| *Katharina tunicata* | SRR11674131 | 15,542 | C:89.7%[S:88.4%,D:1.3%] | Varney, Speiser et al. 2021 |
| *Tonicella lineata* | SRR11577222 | 13,780 | C:79.0%[S:77.7%,D:1.3%] | Varney, Speiser et al. 2021 |
| *Nutallochiton* sp. | SRR11674133 | 57,110 | C:74.3%[S:67.4%,D:6.9%] | Varney, Speiser et al. 2021 |
| *Cryptoplax japonica* | SRR13010086 | 14,963 | C:34.6%[S:34.3%,D:0.3%] | Moles, Cunha et al. 2021 |
| *Cryptoplax larvaeformis* | SRR11674126 | 20,128 | C:88.1%[S:87.7%,D:0.4%] | Varney, Speiser et al. 2021 |
| *Choneplax lata* | SRR13010088 | 16,971 | C:14.3%[S:13.4%,D:0.9%] | Moles, Cunha et al. 2021 |
| *Acanthochitona rubrolineata* | SRP179406 | 44,221 | C:91.8%[S:71.2%,D:20.6%] | ----- |
| *Acanthochitona crinita* | SRR5110525 | 22,678 | C:91.4%[S:91.0%,D:0.4%] | De Oliveira, Wollesen et al. 2016 |
| *Acanthochitona fascicularis* | SRR13862580 | 17,427 | C:88.9%[S:88.5%,D:0.4%] | ----- |

## Material and Methods

### 1. Sample collection

All genomes and transcriptomes used in this study are listed in Table 1. To increase taxon sampling, we newly sequenced an individual of *Schizochiton incisus* and also *Leptochiton asellus. Schizochiton incisus* was collected from a rock on a coral reef at the depth of 80 m of Livock Reef (10°10’N, 115°19’E) by fishing net in the South China Sea on July 11, 2020 (Fig. 1). The whole animals *S. incisus* was preserved in 95% EtOH, which was later stored at room temperature, and a small piece of girdle tissue was removed for DNA extraction. The *S. incisus* sample was deposited in the malacology collections at the Senckenberg Museum, Frankfurt with catalogue number SMF 386201. *Leptochiton asellus* was collected on the rocky shore in September 2019, at Ballyhenry Island, Strangford Lough, at Portaferry, N. Ireland. For *L. asellus*, five tissues, including foot, perinotum, aesthetes, viscera, and shell edge were dissected and fixed in RNAlater (ThermoFisher) at 4 degree and transferred to −80 deep freezer for storage.

**Figure 1.**
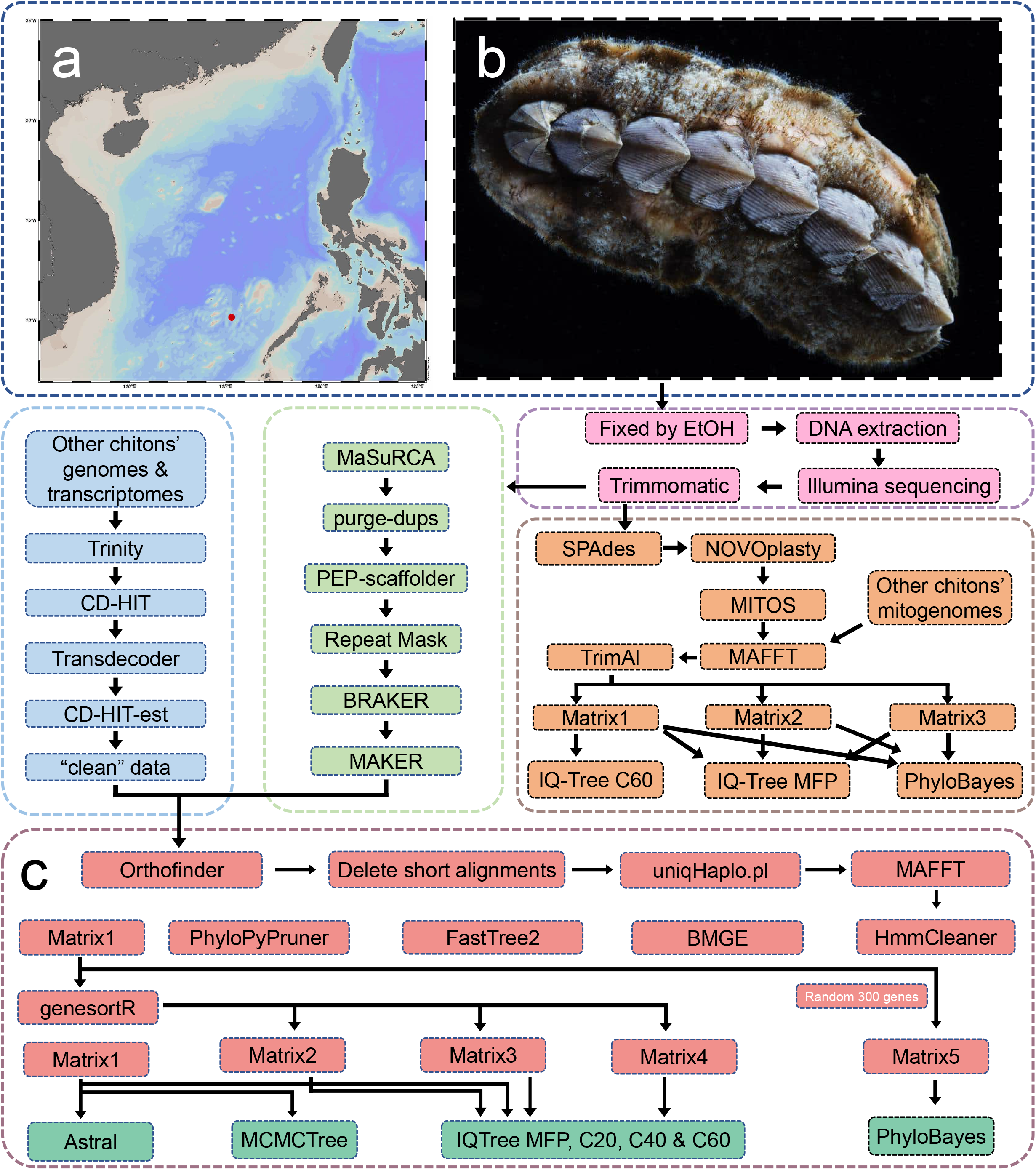
*Schizochiton incisus* (b) and the position where it was collected (a, marked with a red spot). C shows the whole genomic pipeline used in this study, including sample preparation, mitogenome analysis, draft genome assembling and annotation and phylogenomic approach. Photos courtesy of Prof. Xiaoqi Zheng.

### 2. Genome and RNA sequencing

Total genomic DNA of *S. incisus* was extracted with a DNeasy Blood & Tissue Kit (QIAGEN, Germantown, Maryland), which was further sequenced for 150bp paired-end Illumina sequencing to generate approximately 40Gb of raw data on NovaSeq 6000 platform at Novogene (Beijing).

RNA of *Leptochiton asellus* was extracted using Trizol (ThermoFisher) and sent to Novogene (Beijing) for Eukaryotic type transcriptome library preparation and further sequenced on NovaSeq 6000 platform. Approximately 6Gb of raw reads were generated for each tissue.

### 3. Mitogenome analysis

#### 3.1 Mitogenome assemble and annotation

The raw data were trimmed using Trimmomatic v.0.39 (Bolger et al. 2014) with strict filtering settings (ILLUMINACLIP: adapters.fa:2:30:10 LEADING:20 TRAILING:20 SLIDINGWINDOW:4:20 MINLEN:140) to remove low-quality reads and adapters contaminated reads. The resultant clean reads were initially assembled by SPAdes v.3.15.3 (Prjibelski et al. 2020) with default settings, and then the partial COI sequence of *S. incisus* was extracted from the assembled contigs, which was later used as the “seed input” in NOVOplasty v.4.2 (Dierckxsens et al. 2016) to obtain the complete mitogenome of *S. incisus*. The mitogenome was then annotated using the MITOS web server (Donath et al. 2019) with the invertebrate genetic code and the rest default settings, followed by a manual mitogenome annotation confirmation by comparing with other chiton mitogenomes (Irisarri et al. 2020).

#### 3.2 Matrix construction

All *.gb* files of chiton mitogenomes available on NCBI were downloaded and imported into Phylosuite v. 1.2.2 (Zhang et al. 2020), which is an application that allows users to perform phylogenetic analyses on relatively small datasets. All procedures of mitogenome phylogenetic analyses, except for tree constructing and visualization, were carried out through Phylosuite built-in plugins. In brief, 13 protein-coding genes and 2 rRNA genes were extracted from the chiton mitogenomes. Afterwards, MAFFT v. 7.471 was used to align sequences, followed by trimAL v. 1.2rev57 with the “automated1” option to remove spurious sequences and misaligned regions. After that, trimmed sequences were concatenated, generating 3 different matrices. Amino acid sequences of 13 protein-coding genes (PCGs) were extracted and concatenated into a Matrix1. As for Matrix2, all nucleotides of 13 PCGs and 2 rRNA were concatenated. To avoid the phylogenetic signal saturation on the third codon, the third codons of 13 PCGs were replaced by degenerate bases (A, G replaced by R and C, T replaced by Y), then these modified sequences were concatenated, named Matrix3. Generated gene matrix and the corresponding partition file were later used for maximum likelihood (ML) and Bayesian inference (BI) tree construction.

#### 3.3 Mitogenome phylogeny

For the ML framework, IQ-Tree v.2.1.3 (Minh et al. 2020) was implemented using -MFP to select the best-fit model for each partition. Besides, an additional empirical profile mixture model, C60, was also carried out on the AA matrix (Matrix1). All ML analysis were performed with 1000 replicates of ultrafast bootstrapping (-bb 1000).

BI was carried out using PhyloBayes MPI v.1.8c (Lartillot et al. 2013) with CAT-GTR+Γ4 models. For each matrix, four independent Monte Carlo Markov chains (MCMC) were run simultaneously and convergence was checked with the bpcomp program. Then a consensus tree was obtained after discarding the first 10% cycles as a burn-in.

All trees obtained were then visualized with Figtree (http://tree.bio.ed.ac.uk/software/figtree/).

### 4. Genome assembly and annotation

The Illumina raw data was filtered with Trimmomatic v.0.39 (Bolger et al. 2014) with settings of “PE ILLUMINACLIP:TruSeq3-PE.fa:2:30:10 LEADING:10 TRAILING:10 SLIDINGWINDOW:4:15 MINLEN:40”. Afterwards, genome features were calculated by using jellyfish v.2.3.0 (Marçais et al. 2011) (19mer) and GenomeScope2 (Vurture et al. 2017). A benchmark of commonly used assemblers for Illumina data, including Platanus v. 1.2.4 (Kajitani et al. 2014) and MaSuRCA v.4.0.3 (Zimin et al. 2013), was performed based on BUSCO v.5.1.2 score by searching against metazoan odb10 database. Afterwards, purge-dups v. 1.2.5 (Guan et al. 2020) was used to remove redundant contigs, and the resultant contigs were further scaffolded by using PEP-scaffolder (Zhu et al. 2016) with the help of protein sequences from the concatenation of the genome of *Acanthopleura granulata* (Varney et al. 2021).

A custom repeat library of *S. incisus* was *de novo* generated by RepeatModeler v.2.0.2a (Flynn et al. 2020). RepeatMasker v.4.1.0 (Tarailo-Graovac et al. 2009) was performed with the species-specific repeat library mentioned above, followed by a second round of RepeatMasker but with Repbase library 2018 (https://www.girinst.org/repbase/). Afterwards, BRAKER v.2.1.6 (Hoff et al. 2019) was run to train an *ab initio* gene predictor Augustus v.3.4.0 (Stanke et al. 2006) with ODB10 v.1 database downloaded from OrthoDB (Kriventseva et al. 2018), generating a config file of *S. incisus*, which was used as one piece of evidence while running the genome annotator MAKER v.3.01.04 (Holt et al. 2011). Because there was no transcript evidence available, all Mollusca proteins on NCBI were downloaded (Date: Jan 20 2022), and redundancy was removed with CD-HIT v.4.8.1 with the setting of “-c 0.9”. These protein sequences were regarded as the protein homology evidence in MAKER. And the proteins generated from MAKER was used for further phylogenetic analyses.

### 5. Transcriptome assembly and filtration

The protein coding genes of *Acanthopleura granulata, A. loochooana* (Liu et al. 2022) and all other available transcriptomes were downloaded from NCBI SRA database. For transcriptome SRA datasets as well as the transcriptome sequencing of *L. asellus*, the raw reads were *de novo* assembled in Trinity v.2.13.2 or v.2.14.0 (Haas et al. 2013), using the “--trimmomatic” setting, followed by one round of CD-HIT v.4.8.1 (Fu et al. 2012) with the strictest threshold (-c 0.8) to remove redundant sequences. CD-HIT was run multiple times which was continuously monitored by BUSCO5 aiming to get a best score with highest “S” score and lowest “D” (duplicated BUSCO) score. Afterwards, Transdecoder v.5.5.0 (Douglas 2018) was performed to search for open reading frames with the “--single_best_only” option. And the generated peptide files were filtered using CD-HIT with the “-c 0.8” option again to make sure the “D” score wouldn’t drop any more. This step aimed to remove as many heterozygous and transcript isoforms as possible so that they would not mislead orthology inference.

### 6. Orthology inference and matrix construction

Orthology inference was accomplished with a pipeline that was generated from former studies (Kocot et al. 2017; Sun et al. 2021) with slight modifications. We ran Orthofinder v.2.5.4 (Emms et al. 2019) to search for orthologues within selected taxa. Then in the “Orthogroup_Sequences” directory of the Orthofinder output, OG heads were fixed with a custom shell script to make sure that the orthology inference pipeline could be error less. After the preparation, PREQUAL v.1.02 (Whelan et al. 2018) was used to detect and mask non-homologous characters. Then sequences shorter than 100 amino acids were deleted. Occupancy was set to 50%, and redundant sequences were then removed with another custom shell script named uniqHaplo.pl. The leftover *fasta* files were aligned using MAFFT v.7.490 (Katoh et al. 2013) with default settings. Afterwards, HmmCleaner (Di Franco et al. 2019) was used to remove misaligned regions, followed by trimming alignment with BMGE v.1.12 (Criscuolo et al. 2010). Then FastTree2 (Price et al. 2010) was used to construct fast-ML trees for each remaining OGs. Last but not least, PhyloPyPruner v. 1.2.4 (https://pypi.org/project/phylo-pypruner) was performed to identify putative orthology sequences based on the former FastTree2 result, resulting in an initial matrix containing 3593 OGs.

We performed genesortR (Mongiardino Koch 2021) to sort and select “best” OGs based on seven commonly used phylogenetic gene properties, thus genes with best phylogenetic signals can be used for down streaming analysis. An ML tree for the initial matrix was constructed with the IQ-Tree “-MFP” model as input. Also, ML trees for each gene were constructed in IQ-Tree with the same settings. At last, four matrices, including an initial matrix (Matrix1), best 800 genes matrix (Matrix2), best 1300 genes matrix (Matrix3), and best 2700 genes matrix (Matrix4) generated by genesortR, were prepared for phylogenetic analysis.

### 7. Phylogenomics

ML phylogenetic analysis was performed using IQ-Tree 2 (Minh et al. 2020) on the four matrices generated above. The ML approach was carried out using the best-fitting model for each partition (-m MFP). Regarding the *.contree* file generated by the MFP model as the guide tree, PMSF model was then performed in IQ-Tree 2 with site-specific frequency models (C20, C40 and C60). All ML analyses were carried out with 1000 ultrafast bootstrap. As for BI analysis, all matrices mentioned above were too large to run in PhyloBayes MPI v.1.8c, thus the fifth matrix, produced by random 300 genes from Matrix1, was brought out. Four independent chains were run simultaneously until convergent with CAT-GTR+Γ4 model.

A coalescent approach, in contrast to concatenated-based phylogenetic analysis, was also performed to evaluate evolutionary relationships in polyplacophora with ASTRAL v.5.7.1 (Sayyari et al. 2016). An AU-test was performed with IQ-tree 2 on two topologies, which were ((Chitonoidea, Schizochitonoidea), Acanthochitonina) and ((Acanthochitonina, Schizochitonoidea), Chitonoidea), respectively.

## Results

### Mitochondrial genome

We assembled the complete mitochondrial genome of *S. incisus*, which was 15,491 bp in length circularized with 13 PCGs, 2 rRNA, and 22 tRNA, a typical mitogenome architecture of bilaterians. Protein-coding genes are coded with normal invertebrate mitochondrial codons including the start and stop codons. The mitogenome of *S. incisus* follows the proposed hypothetical ancestral gene order for Polyplacophora (Irisarri et al. 2020), except for an inversion of trnG-trnE (Fig. 2b). The mitogenome gene order seems to be relatively conserved in Polyplacophora compared to those in gastropods or bivalves (Irisarri et al. 2020).

**Figure 2.**
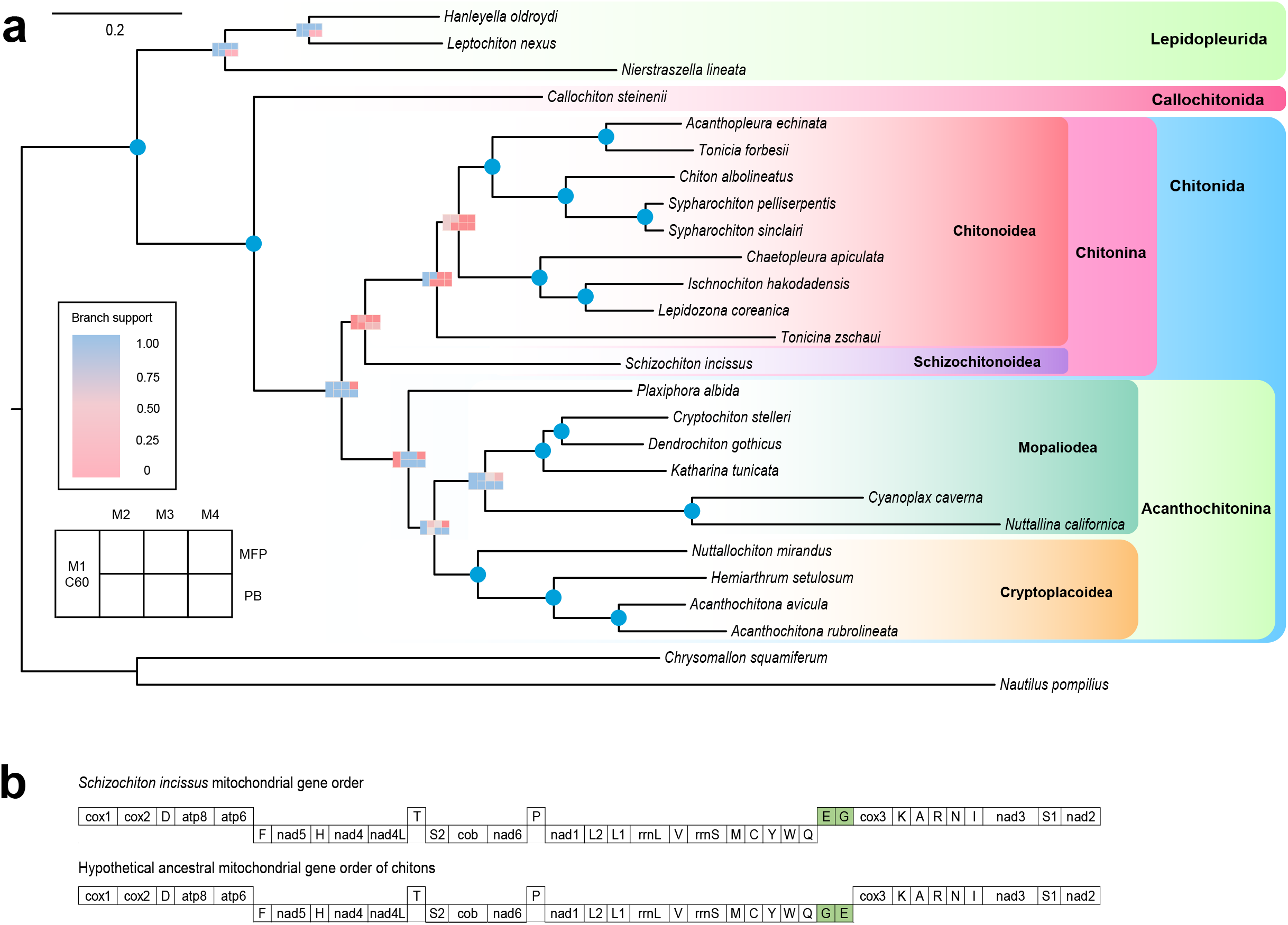
Mitogenome analyses of *Schizochiton incisus*; (a) mitogenome phylogeny of Polyplacophora, and (b) *S. incisus* mitochondrial gene order comparing with hypothetical ancestral mitochondrial gene order of chitons.

### Mitochondrial phylogeny

The phylogenetic trees reconstructed with mitogenome data showed significant discordance among different methods and matrices. There were 3 distinct topologies for the position of *S. incisus*, which were ((Chitonoidea, Schizochitonoidea), Acanthochitonina)(13PCGs with MFP, PB based on modified 3rd codon), ((Acanthochitonina, Schizochitonoidea), Chitonoidea) (13PCGs with C60, PB, PCGs + rRNA with MFP, PCGs + rRNA with PB) and ((Chitonoidea, Acanthochitonina), Schizochitonoidea) (modified 3rd codon), respectively. The statistical support of the *S. incisus* node was lower than 95% in all methods, except for BI, indicating these nodes were not well supported with mitogenomic data. We note that in addition to *Schizochiton*, the position of *Plaxiphora albida* also varied from one clade to another (Fig. 2a). And in the presentative tree, *Tonicina zschaui* was sister to the rest Chitonoidea.

### Genome and transcriptome assembly

Genome features of *S. incisus* were estimated with Illumina sequencing reads, which resulted in an estimated genome size of 1.1 GB and genome heterozygosity of 0.93%. Draft genome assembly from MaSuRCA generated a better result (C:73.8% [S:68.1%, D:5.7%]) than the Platanus version [C:17.9% (S:13.7%, D:4.2%)], which was used for down-stream analyses. After further scaffolding with protein sequences from other chitons with available genomes and removing heterozygous contigs, the final assembly has a BUSCO score of C:73.8%, N50 of 13.2Kb and the assembled size of 971 Mb.

By collecting the evidence from the *ab initial* method and protein evidence, a total of 23,444 protein coding genes were predicted in *S. incisus* with a BUSCO score of C: 40.8% (S: 37.0%, D: 3.8%) and F: 19. 8%. Though the score is lower than the *Acanthochitona rubrolineata* genome (Varney et al. 2021), 12,419 of them (52%) can find their reciprocal best hits BLAST in *A. rubrolineata*, suggesting that a good coverage of protein coding genes for the phylogenomic analyses.

The transcriptome of *Leptochiton asellus* generated from five tissues was assembled into 390,724 contigs with an N50 value of 1.68Kb, and the BUSCO score is C:94.5% (S:83.1%,D:11.4%). For the rest transcriptome assembly of the publicly available data, the BUSCO completeness ranges from 12.9% (*Lepidopleurus cajetanus*) to 95.8% (*Callochiton septemvalvis*) (for the species list and their corresponding BUSCO score, see Table 1).

### Phylogenomics

The phylogenomic analysis was based on the combination of transcriptome and genome data, covering all the extant superfamilies in Polyplacophora (Table 1). There were four matrices generated by genesortR forming seven distinct phylogenetic signals. Minimum occupancy for all matrices was set to 50%. The sites contained in the four matrices are 696,897 (3593 genes, all genes, Matrix 1), 194,356 (best 800 genes, Matrix 2), 299,710 (best 1300 genes, Matrix 3), 554,857 (best 2700 genes, Matrix 4), respectively (Fig. 3).

**Figure 3.**
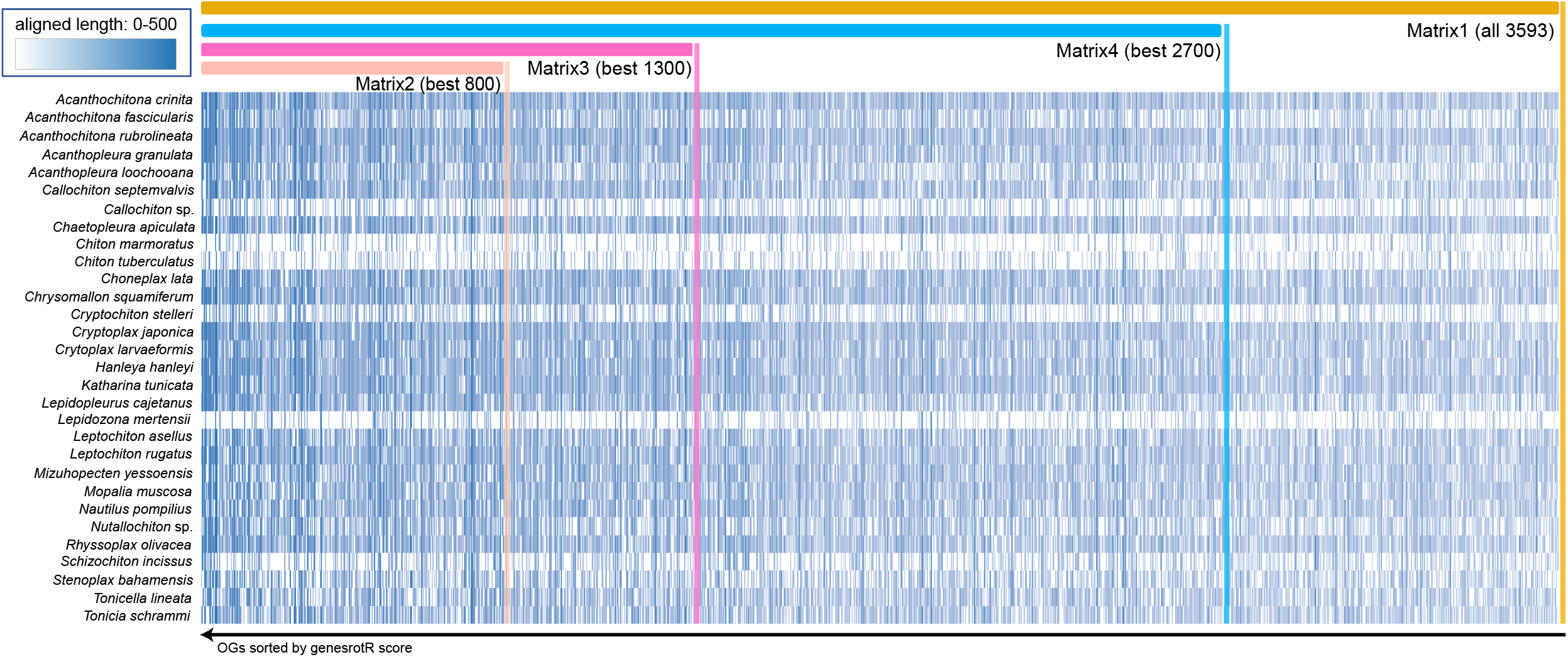
The occupancy of the four matrices generated by genesortR.

The phylogenetic trees reconstructed from nuclear data, including coalescent approach results, showed a high degree of consistency about the position of *S. incisus*, as sister to Chitonoidea (Fig. 4). Support for this Schizochitonoidea + Chitonoidea clade retrieved node support of 100% in all analyses except for PMSF-C20 of Matrix1 (which is 59), showing a relatively stable topology. The support for all superfamily level groups and their arrangement was consistently high. However, the positions of some tips are unsettled: *Chaetopleura apiculata*, *Lepidozona mertensii* and *Stenoplax bahamensis* resolved in variable positions within the superfamilies. The relationship of *Choneplax* relative to the members of genus *Acanthochitona* is also changeable.

**Figure 4.**
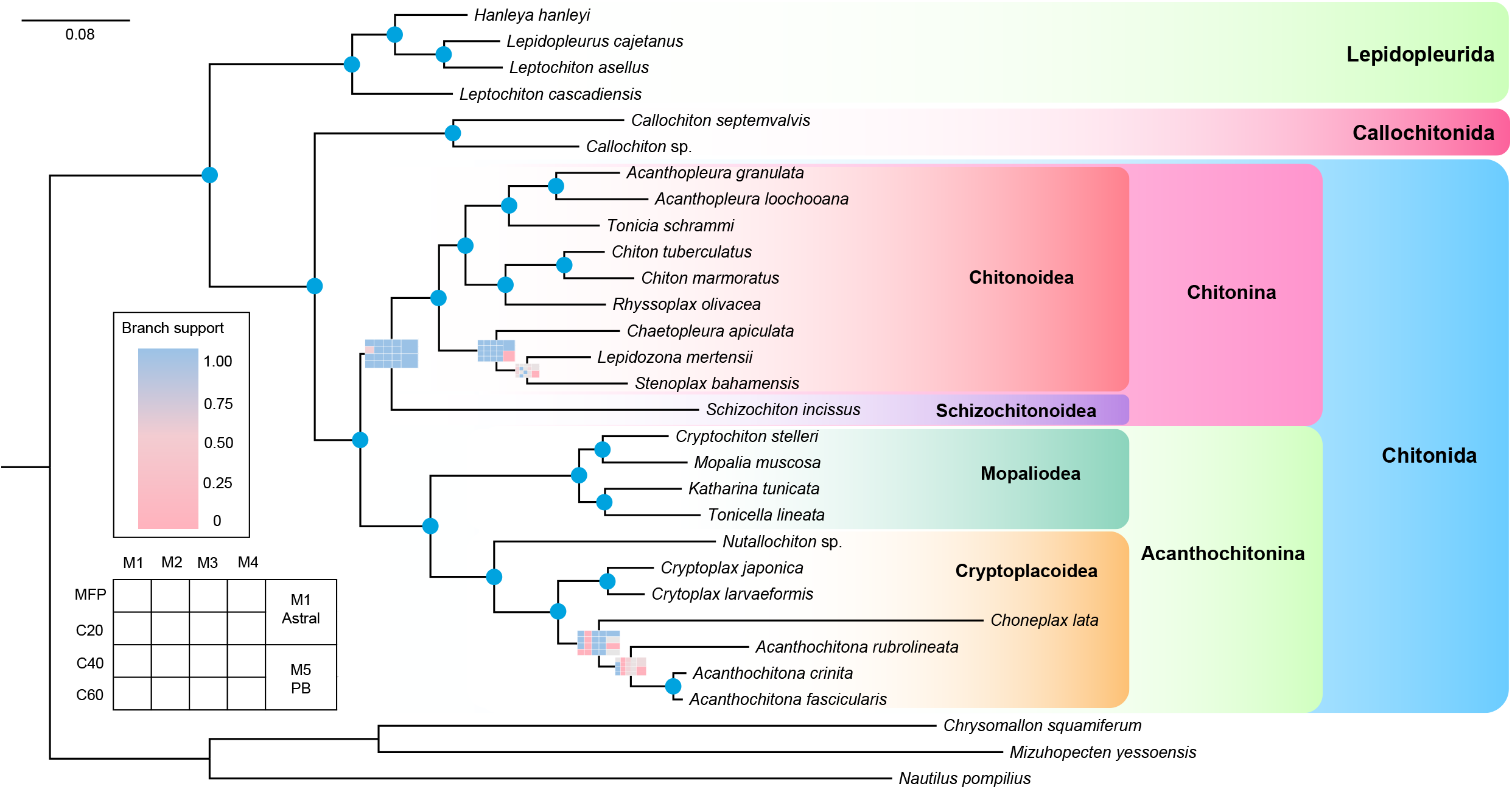
Phylogeny of chiton based on phylogenomic approach with different methods. Node support are transferred into matrices colored with a continuous scale bar ranging from 0 to 1. Blue indicates 100% support and pink indicates the topology is not supported by the representative tree. And node with blue spot indicates full support in all methods. M1-M5, matrix 1-5; MFP, IQ-Tree MFP model; C20-C60, profile mixture models C20-C60; M1 Astral, coalescent analysis based on Matrix1; M5 PB, PhyloBayes analysis based on Matrix5.

### Topology test

We performed AU-test on two topologies based on Matrix1 to determine the better supported tree topology. Given results with *P*-value < 0.05 will be rejected. The results showed that the first tree topology ((Chitonoidea, Schizochitonoidea), Acanthochitonina) was accept with a *P*-value of 0.952, and the second topology ((Acanthochitonina, Schizochitonoidea), Chitonoidea) was rejected with a *P*-value of 0.0476.

## Discussions

The phylogenetic relationships of chiton at the order and superfamily levels are relatively stable and well resolved. Based on a consensus of phylogenetic analyses, Polyplacophora is divided into three orders, Lepidopleurida, Callochitonida and Chitonida (Irisarri et al. 2020; Moles et al. 2021), which is also recovered in the present analyses. At the superfamily level, former molecular studies lacked data to test the position of Schizochitonoidea, and our results support the sister relationship of Chitonoidea + Schizochitonoidea in a monophyletic suborder Chitonina, as proposed from integrated morphological and anatomical evidence (Sirenko 2006).

The mitogenome data were much less informative than nuclear transcriptome and genomic data. We used mitogenome data of available chitons to reconstruct phylogenetic trees with different approaches, including ML and Bayesian inference, but the results below superfamily level are unstable. For example, in the representative tree selected for mitochondrial analyses, *Tonicina zschaui* formed a sister group to other remaining Chitonoidea, whereas current systematics would predict a placement for *Tonicina* within the small clade formed by the genera *Lepidozona, Ischnochiton* and *Chaetopleura*. The topology we illustrated is not supported by 4 of 7 trees reconstructed by corresponding methods, so this placement should be taken as unresolved. As already suggested in the previous mitogenome phylogeny of chitons (Irisarri et al. 2020), this could be a result by poor taxon sampling, but it was not improved by adding a few additional taxa here. Indeed, this issue of low phylogenetic signal in mitogenome phylogeny has also been raised in data from another molluscan class, Monoplacophora (Stoger et al. 2016), and confounding features occur in many molluscan mitogenomes (Ghiselli et al. 2021).

Interestingly, *Schizochiton* possesses a unique mitogenome gene order, differing from any other chitons with available mitogenomes, which might imply relatively fast evolution of the species. Mitogenome phylogenies are currently not reliable for reconstructing detailed phylogenies for Polyplacophora and potentially other molluscan clades. This may be improved with better taxon sampling, or may be a fundamental problem of insufficient phylogenetic signal. It is clear that currently phylogenomic approaches are needed to reconstruct phylogeny of chitons at or below superfamily level resolution.

All the phylogenomic results for the main lineage in this study shared the same topology with strong node support except for Matrix1-C20. The topology is consistent with what is by now a well-established backbone phylogeny for Polyplacophora and also concordant at superfamily and higher level with the mitogenome phylogeny (Sigwart et al. 2013; Irisarri et al. 2020; Moles et al. 2021). Lepidopleurida is sister to the remaining Polyplacophora. *Callochiton*, representing the order Callochitonida is sister to Chitonida. This latter order is divided into two clear clades representing the suborders Chitonina and Acanthochitonina.

Our phylogeny of polyplacophora based on phylogenomic approach possesses more advantages than former molecular studies (Okusu et al. 2003; Sigwart et al. 2013; Irisarri et al. 2020; Moles et al. 2021), including a broader taxon sampling and massive genes, has resolved the relationships among main lineages of chitons. The genus and family level arrangement of taxa in this study are largely concordant with established taxonomy or with other molecular studies from smaller data matrices. Within Lepidopleurida, the family

Leptochitonidae s.s. is restricted to the NE Atlantic species, represented here by *Leptochiton asellus* and *Lepidopleurus cajetaus*, with the Pacific *Leptochiton cascadiensis* outside that clade, as is already established from previous molecular studies using Sanger sequencing (Sigwart et al. 2011; Sigwart 2016).

Acanthochitonina is known to be divided into two clades based on egg hulls and hexagon edges projections; one of the clades, Mopalioidea, includes *Cryptochiton*, *Mopalia*, *Katharina* and *Tonicella* (Okusu et al. 2003) which is also well supported by every other molecular phylogeny including our results and modern phylogenetic systematics (Sigwart et al. 2013). Family level arrangement is difficult to test with limited taxon sampling, but the genera in our study group into these four genera that are closely allied to Mopaliidae as separate from a second clade of *Nuttallina* + *Cyanoplax*, also as found in previous studies (Irisarri et al. 2020). In the other superfamily Cryptoplacoidea, *Nuttallochiton* is sister to the rest of Cryptoplacoidea, in accordance with previous molecular studies (Okusu et al. 2003; Sigwart et al. 2013; Irisarri et al. 2020). However, the position of *Plaxiphora* within Acanthochitonia is equivocal; this has been a persistent problem in every molecular phylogeny of chitons, although multiple morphological characters unite *Plaxiphora* with the family Mopaliidae (Sirenko 2006).

*Schizochiton* resolved as sister to Chitonoidea, forming a monophyletic suborder Chitonina with full support except for Matrix1-C20 method, being sister group with the larger order Chitonida. The only prior molecular analysis to include *Schizochiton* also recovered it as sister to the remaining Chitonina in one version of their analyses, but concluded that its position within the phylogeny was effectively unresolved (Okusu et al. 2003: fig 5). The position of *Schizochiton* was controversial because of an unusual combination of morphological characters. The balance of evidence placed this group in the suborder Chitonina (Sirenko 2006). *Schizochiton* possess a caudal sinus in tail valve that is similar to others in Mopalioidea as well as egg hulls with cupules that are simpler but comparable to other Mopalioidea. Based on the new phylogenetic tree, we can infer these features may be plesiomorphic for the larger order Chitonida.

One important morphological feature of *Schizochiton* that differs from almost all other chitons is their shell eyes. Shell eyes were described in 1884 from specimens of *Schizochiton incissus* (Moseley 1884), and were immediately recognized as modifications of the chiton aesthete system (Moseley 1885). All chitons possess aesthete pores in their shell plates and some are photosensitive (Kingston et al. 2018). But shells eyes are restricted to only a few genera, in the family Schizochitonidae and the family Chitonidae. Those genera in the family Chitonidae with shell eyes form a monophyletic clade and have a fossil record only dating back to the Miocene (Sirenko 2006). Phylogenetic and fossil evidence suggests that shell eyes evolved first in Schizochitonidae and again a second time very recently in the history of Chitonidae.

Recognizing *Schizochiton* within a superfamily level group Schizochitonoidea, sister to Chitonoidea, confirms the relationship predicted by morphological systematics. This is now confirmed from molecular evidence and a more stable phylogeny than earlier preliminary results. This also reaffirms that the multiple lines of evidence from morphological, anatomical, and gamete characters already recognized in chitons provide a robust basis for phylogenetic systematics.

## Data availability

The raw Illumina sequencing data was deposited on NCBI SRA database with the accession No. of PRJNA909482, and the assembled mitogenome on NCBI nucleotide database with the No. of XXXX. The assembled genomic contigs, predicted gene models can be accessed via FigShare with the URL of 10.6084/m9.figshare.21709742.

## Acknowledgements

This research project was financially supported by the Fundamental Research Funds for the Central Universities (202241002 and 202172002), Science and Technology Innovation Project of Laoshan Laboratory (No. LSKJ202203100), and the Young Taishan Scholars Program of Shandong Province (tsqn202103036). Bioinformatic analysis was conducted on the high-performance server IEMB-1 hosted at Institute of Evolution and Marine Biodiversity. We also thank Dr. Chong Chen (JAMSTEC) for help with fieldwork and specimens. This is contribution number 14 from the Senckenberg Ocean Species Alliance.

